# A biomimetic electrical stimulation strategy to induce asynchronous stochastic neural activity

**DOI:** 10.1101/725432

**Authors:** Emanuele Formento, Edoardo D’Anna, Sandra Gribi, Stéphanie P. Lacour, Silvestro Micera

## Abstract

Electrical stimulation is an effective method to communicate with the nervous system. Here, we demonstrate that current stimulation paradigms fail to reproduce the stochastic and asynchronous properties of natural patterns of neural activity, and introduce a novel biomimetic stimulation (BioS) strategy that overcomes these limitations. We hypothesized that high-frequency amplitude-modulated bursts of stimulation could induce asynchronous neural firings by distributing recruitment over the duration of a burst, without sacrificing the ability to precisely control neural activity. We tested this hypothesis using computer simulations and ex vivo experiments. We found that BioS bursts induce asynchronous, stochastic, yet controllable, neural activity. We established that varying the amplitude, duration, and repetition frequency of a BioS burst enables graded modulation of the number of recruited fibers, their firing rate, and the synchronicity of the responses. These results demonstrate an unprecedented level of control over artificially induced neural activity, enabling the design of next-generation biomimetic stimulation paradigms with potentially profound consequences for the field of neurostimulation.

## Introduction

Electrical stimulation is a safe and effective method to interact with the nervous system, which has been successfully used in neuroprosthetics to treat brain disorders^1–3^, control cardiac activity^4^, alleviate chronic pain^5^, restore lost sensory and motor functions^6–11^, and even control autonomic functions^12^. During electrical stimulation, electric currents are exchanged between a stimulator and a pair of electrodes in contact with the body. This process involves the accumulation of charges and electrochemical reactions (oxidation and reduction of ionic species) at the electrode/tissue interface, as reviewed in detail by Merrill and colleagues^13^. Most commonly, charge-balanced square biphasic pulses are used to activate neural structures, whereby a cathodic current-controlled square pulse is delivered first, followed by an anodic pulse of equal amplitude. This design achieves a good tradeoff between effectiveness and safety^13,14^. Single biphasic pulses represent the fundamental units of stimulation, but are generally delivered in trains to modulate neural circuit dynamics over time. In this manuscript, we refer to this common electrical stimulation paradigm as “pulsed stimulation”, whereby a train of cathodic-first biphasic pulses is delivered at frequencies lower than the maximal firing rate of the target neural structures.

Three parameters are used to adjust the effects of stimulation, namely stimulation amplitude, pulse width, and frequency. Both amplitude and pulse width control the size of the population of neurons recruited by each pulse of stimulation, while stimulation frequency controls the rate at which neurons are recruited. These parameters are tuned on an application-specific basis^8,15^, often with the goal to recruit neural structures in a way that most closely matches their natural idiosyncratic activity^8,9,16–19^. However, pulsed stimulation recruits entire neural populations in synchrony with each stimulation pulse^20^, with a maximum spread dictated by the pulse width, which typically ranges from a few hundreds of microseconds to a few milliseconds at most. Such synchronous activity is not commonly observed in-vivo, where neural activity is often largely asynchronous, driven in part by the probabilistic nature of action potential generation in sensory organs, such as muscle spindles^21^ or retinal cells^22^, and in part by the stochastic nature of synaptic transmission^23^.

The high synchronicity of neural activity resulting from pulsed stimulation can have negative consequences in several neurostimulation applications. For instance, in the context of tactile feedback restoration for amputees, it has been hypothesized that the synchronous neural activity induced in populations of cutaneous afferents accounts for the difficulty in producing natural tactile percepts^16,20^. Similarly, in the case of functional electrical stimulation (FES), where stimulation is used to induce muscle contractions, the synchronicity of the response has been shown to induce more jerky movements and higher muscle fatigue^9,24,25^.

Generating biomimetic, asynchronous neural activity with electrical stimulation remains an unsolved challenge. A recent study proposed replacing the cathodic and anodic phases of each stimulation pulse with a high-frequency burst of short pulses (approximately 100 *μ*s, delivered at more than 7 kHz), with a longer total duration^26^. Using this approach, Zheng et al. showed that it is possible to spread motor-axon recruitment over a window of approximately 2 ms, thus inducing motor responses that are somewhat less synchronous compared to pulsed stimulation (see Table 1 of Zheng et al^26^). However, while this study demonstrates a first step towards inducing asynchronous and controllable neural activity, natural firing patterns remain drastically less synchronized, and the proposed approach is unlikely to enable spreading neural activity over longer periods of time, since delaying the delivery of the anodic phase for too long will increase the chances of tissue damage^13^.

Here, we first studied how neural activity induced during pulsed stimulation differs from that generated during natural stochastic processes. Using computer simulations, we emphasized that electrically-induced activity is not only more synchronous, but also less variable. We then proposed a biomimetic stimulation (BioS) strategy in which each individual stimulation pulse is replaced by a burst of amplitude-modulated high-frequency stimulation. We reasoned that by slowly increasing the stimulation amplitude, a burst of high-frequency pulses would first recruit the most excitable neurons (large-diameter fibers and neurons near the electrode), and then progressively recruit less excitable cells. In addition, we hypothesized that by using high-frequency stimulation (in the kHz range), the recruited neurons would rarely fire more than once per burst, thus allowing the distribution of neural recruitment over indefinitely long bursts, while still being able to control the number of action potentials. Indeed, high-frequency stimulation waveforms have been shown to elicit only transient neural activity, with neurons firing only once to a few times at the stimulation onset and then becoming unresponsive to further stimulation pulses until the stimulation is either stopped or restarted^27–30^. Combining computer simulations and ex vivo experiments in nerve fibers, we corroborated these hypotheses, and showed that this strategy allows for the generation of asynchronous, stochastic, yet controllable, neural activity. We argue that the ability to desynchronize neural activity during electrical stimulation could have far-reaching implications in several applications of neurostimulation.

## Results

### Pulsed stimulation induces synchronous and regular neural activity

We investigated the ability of pulsed electrical stimulation to induce biomimetic patterns of neural activity in a population of myelinated fibers. We first developed a biophysical model of a bundle of fibers—composed of a heterogeneous population of group-II fibers, 40 cm in length in an isotropic medium—and evaluated the effect of electrical stimulation on the fibers’ membrane potential (Figure 1a and b). We simulated a train of biphasic, charge-balanced, symmetric stimulation pulses at an amplitude of 0.8 mA and a frequency of 40 Hz (Figure 1c). Each stimulation pulse instantaneously induced an action potential in the majority of the modeled fibers (71%). Action potential generation occurred at the node closest to the electrode. Since conduction velocity depends on fiber diameter, the heterogenous distribution of diameters (9.2 ± 0.5 μm) within the modeled fiber population caused a weak, progressive desynchronization as the action potentials traveled away from the site of stimulation. After travelling along the entire length of the fiber, the time difference between the earliest and the latest action potentials reached approximately 3 ms (Figure 1c).

**Figure 1.**
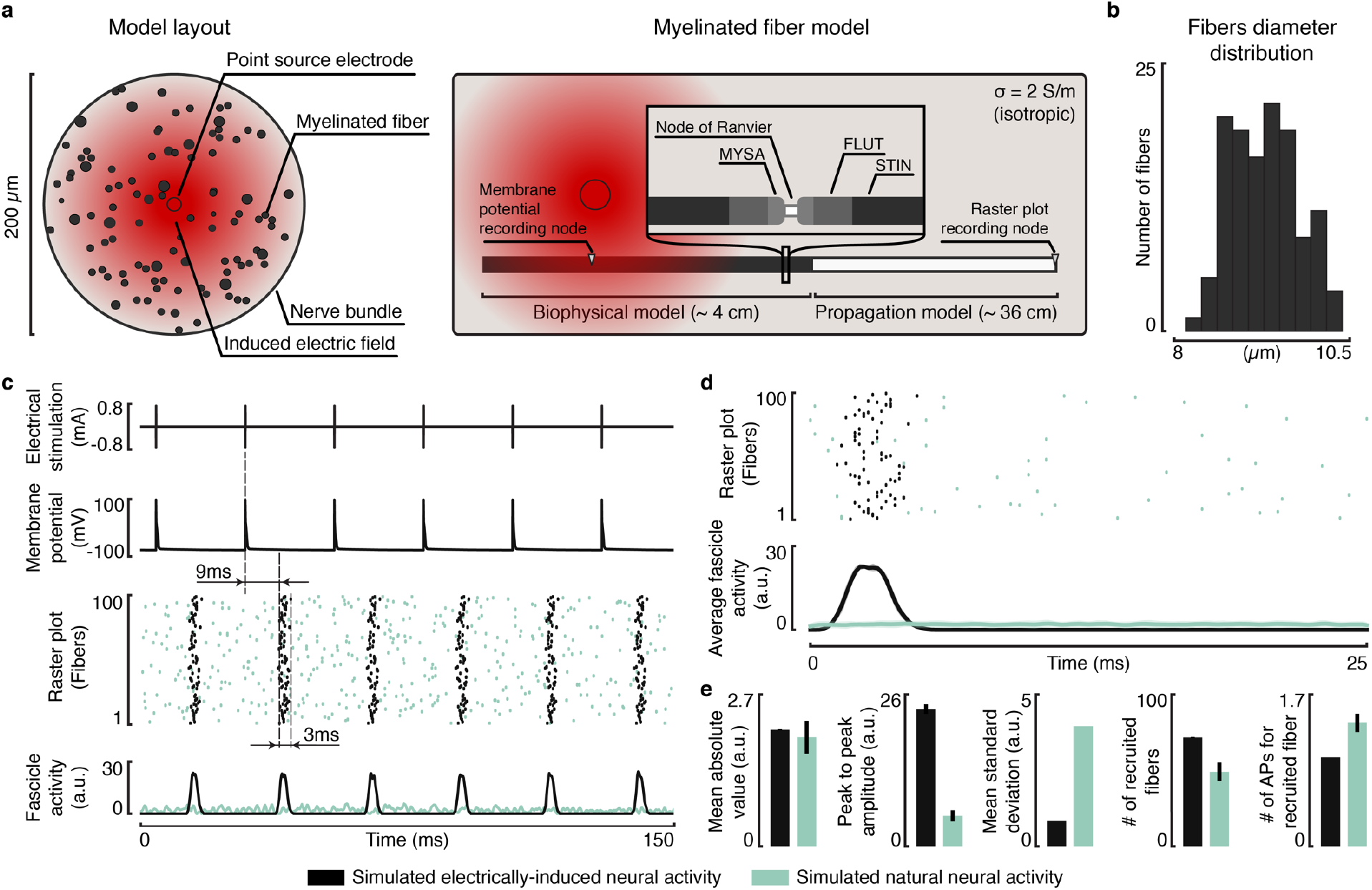
Pulsed electrical stimulation induces synchronous and regular neural activity: a modeling study. **a**, Layout of the biophysical model used to evaluate the effect of electrical stimulation on a population of myelinated fibers. Each fiber is composed of two components: a multicompartmental model that describes the physics of the fiber’s nodes of Ranvier and of the myelinated segments, which we used to evaluate the effect of an external electrical field on the fiber potential; and a propagation model that simulates the propagation of the action potentials generated in the multicompartmental model along the whole length of the fiber (40 cm). We modeled the effect of electrical stimulation by considering a point source electrode located in the center of the population, and assuming an infinite, homogeneous, isotropic space. **b**, Distribution of the modeled fibers’ diameter. **c**, Effect of pulsed stimulation (40 Hz, 0.8 mA, and 50 μs pulse width) on the modeled population, and comparison with simulated natural firings. The panels report, from top to bottom: the simulated electrical stimulation profile; the effect of electrical stimulation on the membrane potential of a representative fiber at the closest node to the electrode; a raster plot reporting the induced spiking activity reaching the end of the modeled fibers (black dots); and the estimated population activity (black curve). The aquamarine dots and curve represent the simulated natural spiking and population activity, respectively. **d**, Representative spiking activity induced electrically (black) or naturally (aquamarine) during a binned stimulation period (top), and average (± standard deviation) population activity over all the simulated stimulation periods (bottom). **e**, Bar plots representing the mean and standard deviation of the statistics used to compare the electrically induced activity from the natural one, which we computed over the activity binned in stimulation intervals (n = 18 responses, for each condition). From left to right, the statistics are: the mean absolute value, which we adopted as a measure of the overall level of neural activity; the peak to peak amplitude, measuring the synchronicity of the neural activity (synchronous responses have higher peak to peak amplitudes); the mean standard deviation of the neural responses (no standard deviation is reported); the number of recruited fibers; and the average number of action potentials (APs) induced in the recruited fibers.

We next evaluated the difference between electrically-induced and natural neural responses at a given mean firing rate (Figure 1c and d), simulated using Poisson processes. We estimated nerve activity during electrical stimulation and during natural firing by convolving each action potential with a Gaussian wavelet (Figure 1c and d). We then binned the extracted activity into stimulation intervals (25 ms, Figure 1d) and computed three statistics: the mean absolute value, as a measure of the overall level of neural activity; the peak-to-peak amplitude, as a measure of the synchronicity of the response (since summation of multiple synchronous action potentials causes higher maximum values); and the mean standard deviation, as a measure of the response variability. In addition, we computed the average number of action potentials per recruited fiber and the overall number of fibers recruited by the stimulation, to assess fiber recruitment (Figure 1e). Since we imposed the same mean firing rate, the mean absolute value of electrically-induced and natural neural activity was similar (Figure 1e). However, the peak-to-peak amplitude was, on average, 344 % greater during electrical stimulation, suggesting that stimulation-induced neural responses were much more synchronous than natural activity (Figure 1e). Electrically-induced neural responses were also characterized by lower variance (Figure 1e). This was caused by the repeatability of fiber recruitment at each pulse of stimulation, which contrasts with the stochasticity of natural neural activity. These results highlight a sharp discrepancy between naturally occurring neural activity and the neural activity triggered by pulsed electrical stimulation.

### BioS: A stimulation strategy to induce biomimetic patterns of neural activity

Next, we designed an electrical stimulation strategy—named BioS, for **Bio**mimetic **S**timulation— to induce neural patterns that are more similar to naturally occurring activity. This strategy consists in replacing each stimulation pulse with an amplitude-modulated high-frequency (1-10 kHz) burst of charge-balanced biphasic pulses (Figure 2a). Two hypotheses serve as the theoretical basis for this strategy. First, we reasoned that by progressively increasing the stimulation amplitude of each pulse within a burst, fiber recruitment would be distributed over time: large-diameter fibers, or fibers close to the electrode, would be recruited towards the beginning of the burst, as their recruitment threshold is lower; small-diameter fibers, or fibers further from the electrode, would be recruited towards the end of the burst, as their recruitment thresholds is higher (Figure 2a).

**Figure 2.**
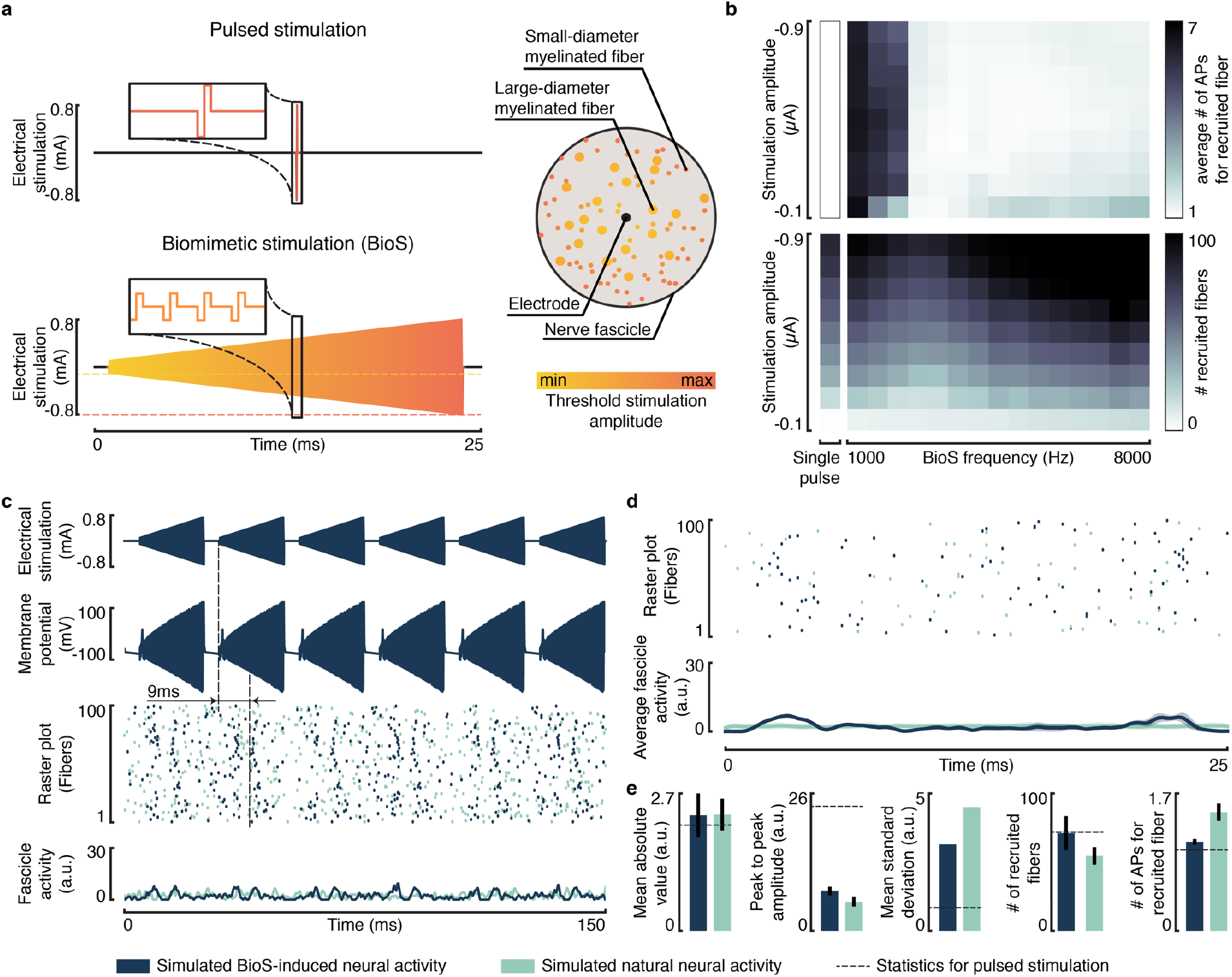
Proposed Biomimetic Stimulation (BioS) strategy. **a**, Fiber recruitment during a BioS burst. BioS amplitude-modulated high-frequency bursts are designed to recruit the same number of fibers as a single stimulation pulse, while distributing fibers’ recruitment throughout their entire duration. Large-diameter fibers, or fibers close to the electrode are recruited at the lowest amplitudes (i.e., at the beginning of the burst), since their stimulation threshold is the lowest. Small-diameter fibers, or fibers far from the electrode are recruited at higher amplitudes (i.e., towards the end of the burst), since their stimulation threshold is higher. **b**, Effect of BioS burst frequency and maximum amplitude on the number of action potentials (APs) generated in each fiber recruited by the stimulation (top), and on the number of recruited fibers itself (bottom). **c**, **d** Effect of 40 Hz BioS (burst frequency: 3500 Hz, burst duration: 20 ms, amplitude: 0.8 mA, pulse width: 50 μs) on the modeled population of fibers, and comparison with simulated natural firings. The same conventions adopted in Figure 1 are used. **e**, Bar plots representing the mean and standard deviation of the statistics used to compare BioS-induced neural activity (navy-blue bars) from both the simulated natural activity (aquamarine bars) and the neural activity induced by pulsed stimulation (dashed lines), as shown in Figure 1e (n = 18 responses, for each condition).

Second, since neuronal membranes cannot follow high-frequency electric field oscillations, but can exhibit transient responses^27–30^, we hypothesized that the recruited fibers would rarely fire more than once per burst. Therefore, each burst would approximately trigger approximately the same number of action potentials within the target population as a single pulse delivered at the highest amplitude within the burst, while desynchronizing the induced activity by recruiting fibers throughout its entire duration. Finally, we hypothesized that adjusting the burst amplitude, duration, and repetition frequency would enable precise control over the induced neural activity.

To evaluate the proposed stimulation strategy, we first used the computational model described above to characterize the effect of different burst frequencies and amplitudes on fiber recruitment (Figure 2b). When using burst frequencies above 2 kHz, simulations indicated that recruited fibers rarely fire more than once. Increasing the maximum burst amplitude had limited influence on the number of times a fiber was recruited, but increased the overall number of recruited fibers (Figure 2b). Interestingly, increasing the frequency above approximately 3.5 kHz also increased the number of recruited fibers. At these frequencies, BioS bursts recruited more fibers than a single stimulation pulse delivered at the same amplitude—a phenomenon caused by the build-up of neuronal membrane depolarizations from individual pulses throughout the burst^31,32^. Since the effects of individual sub-threshold pulses are short lived (due to passive membrane properties), only two stimulation pulses occurring in quick succession can interact with each other to produce facilitation. At lower burst frequencies (<3.5kHz), such interactions did not occur.

We then assessed the similarity between the neural activity induced during three different conditions: a BioS burst repeated at 40 Hz (burst duration: 20 ms, burst frequency: 3.5 kHz), simulated natural neural activity with a mean firing rate set to be equal to the BioS condition, and pulsed stimulation (Figure 2c-e). As hypothesized, BioS-induced neural activity displayed lower synchronization compared to pulsed stimulation, with action potentials distributed throughout the entire burst duration (Figure 2c and d). Indeed, the average peak-to-peak amplitude decreased by 68% with respect to pulsed stimulation, and was only 40% greater compared to natural activity (Figure 2e). In addition, only a minimal increase in mean absolute value was observed. Indeed, although the total number of recruited fibers remained the same as with pulsed stimulation, each recruited fiber fired 1.1 times on average, explaining the small increase in overall neural activity. Finally, BioS bursts induced responses displayed a markedly higher variability compared to pulsed stimulation, with a mean standard deviation approaching the value obtained using simulated natural neural activity (Figure 2e).

These results largely confirm the theoretical foundations of our hypotheses and suggest that BioS stimulation could be used to artificially induce neural activity which is both highly asynchronous and highly variable. These two characteristics are essential for eliciting biomimetic patterns of neural activity.

### BioS bursts induce asynchronous stochastic neural activity

To validate the predictions obtained with our model, we designed an ex vivo experiment using explanted rat dorsal rootlets and a nerve-on-a-chip platform^33^. This platform allowed us to electrically stimulate the small explanted population of heterogeneous fibers (100-150 fibers, Supplementary figure 1) and to measure the induced activity 1.3 cm away from the location of stimulation (Figure 3a and b).

**Figure 3.**
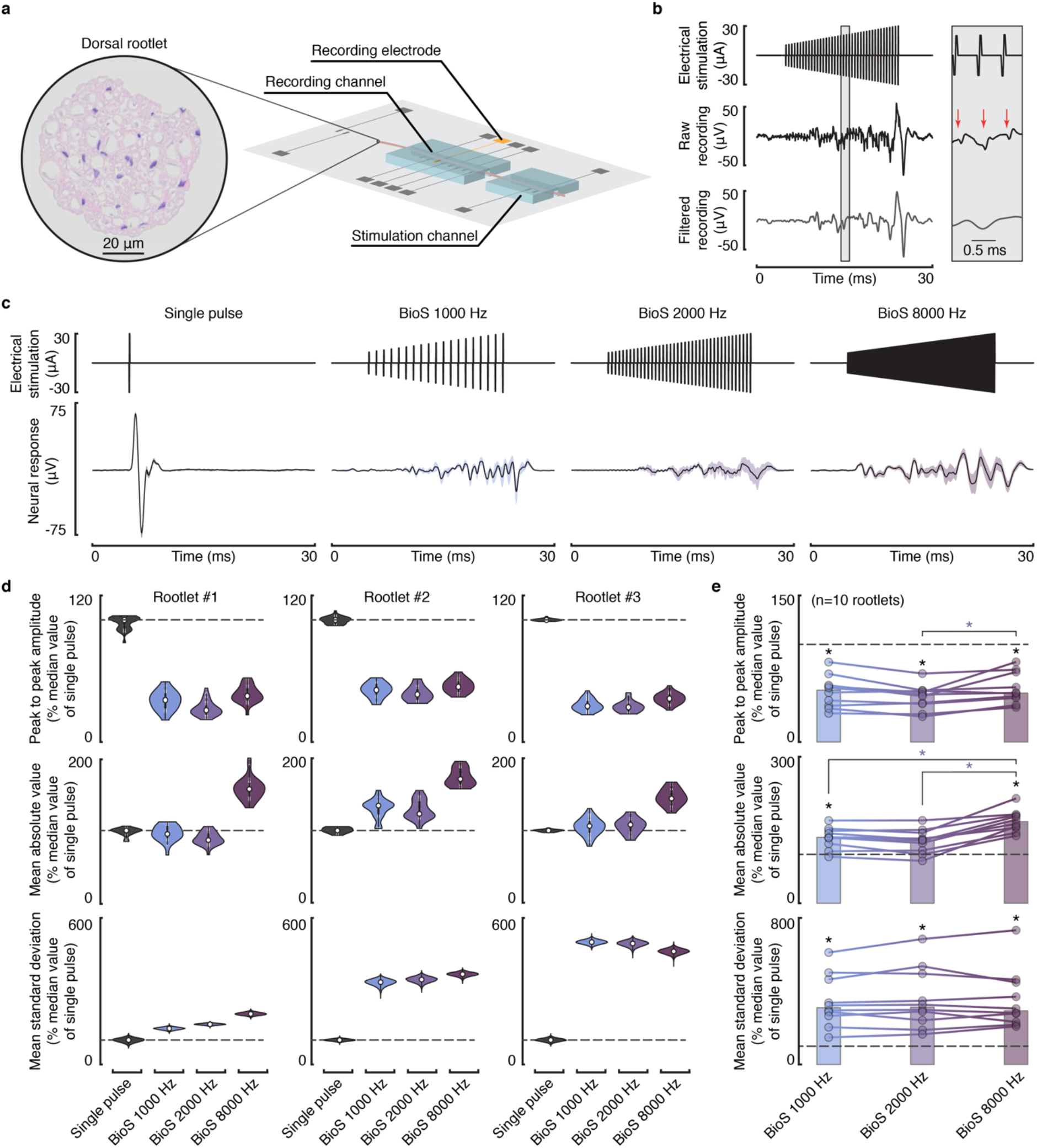
Effect of a BioS burst on heterogeneous populations of rat afferent fibres. **a**, Experimental setup. Extracted rat dorsal rootlets are inserted in a nerve-on-a-chip platform that allows electrical stimulation and subsequent recording of the induced responses a 1.3 cm away from the stimulation site. **b**, Representative recording of the neural response induced by a BioS burst, before and after artefacts removal. The box on the right is a zoom-in of the highlighted time period; red arrows indicate the stimulation artefacts. **c**, Average (± standard deviation) of the neural responses induced by single stimulation pulses, and by BioS bursts delivered at 1000, 2000, and 8000 Hz, respectively from left to right, in a representative rootlet (rootlet #1; burst duration: 20 ms; pulse width: 50 μs). **d**, Violin plots reporting, from top to bottom, the peak to peak amplitude, mean absolute value, and mean standard deviation of the neural responses induced by the tested stimulation conditions, for 3 (over 10) representative rootlets (n = 112, 102, 148 responses, for rootlet #1, #2, and #3, respectively). Each statistic is normalized with respect to the median value of single pulse stimulation, as highlighted by the dashed lines. The violin plots for the peak-to-peak amplitude and the mean absolute value report the sampled distributions. Violin plots for the mean standard deviation report the bootstrapped distribution (n iterations = 10,000). **e** Scatterplots reporting the median of the computed statistics for each BioS condition, for all (n=10) the tested rootlets. For each rootlet, statistics are normalized with respect to the median value of single pulse stimulation, highlighted by the dashed lines. Lines connecting different BioS conditions indicate the results for different rootlets. Bars represent the median of each statistic across the tested rootlets. Black stars indicate a significant difference between the starred BioS condition and 1 (i.e., the median value of single pulse stimulation); p<0.05, two-sided Wilcoxon signed rank test with Bonferroni correction for multiple comparisons. Violet stars indicate a significant difference between two BioS conditions; p<0.05, two-sided, paired Wilcoxon signed rank test with Bonferroni correction for multiple comparisons.

We stimulated 10 dorsal rootlets with either single pulses, or 20 ms long BioS bursts with frequencies of 1000 Hz, 2000 Hz, and 8000 Hz. We used an interval of 2 s between consecutive repetitions. Figure 3c reports the average (± standard deviation) response of one dorsal rootlet to these four stimulation conditions. Single pulses of stimulation activated the entire population of recruited fibers simultaneously, producing a large compound action potential in the measured signal. In contrast, BioS bursts recruited fibers throughout their entire 20 ms duration, without inducing an initial synchronized response. To analyze these responses, we computed the same statistics we adopted in our simulations, namely peak-to-peak amplitude, mean absolute value, and mean standard deviation (Figure 3d and e, and Supplementary Figure 2). Results across the 10 tested rootlets show that the peak to peak amplitudes of the responses obtained with BioS bursts were significantly lower (median 47% lower) than those obtained with single pulse stimulation, for all tested burst frequencies (Figure 3e, * p<0.05, two-sided Wilcoxon signed rank test with Bonferroni correction for multiple comparison). Regarding mean absolute value, we found no significant difference between the responses of BioS bursts delivered at 2000 Hz and those induced by single pulses of stimulation (yet, the median was 29% higher). However, BioS bursts delivered at 1000 Hz and 8000 Hz led to responses with significantly higher mean absolute value. Finally, BioS bursts induced responses with drastically higher mean standard deviations (Figure 3e).

These results validate our model predictions, suggesting that each BioS burst induces asynchronous and stochastic neural activity, rarely recruiting individual fibers more than once (especially at 2000 Hz).

### BioS burst amplitude-modulation is essential for desynchronizing neural responses

Next, we evaluated whether amplitude-modulation of a high frequency burst is necessary to desynchronize neural activity. We stimulated the extracted dorsal rootlets with constant-amplitude high-frequency bursts of 20 ms at 1000 Hz, 2000 Hz, and 8000 Hz, and evaluated the neural responses (Figure 4).

**Figure 4.**
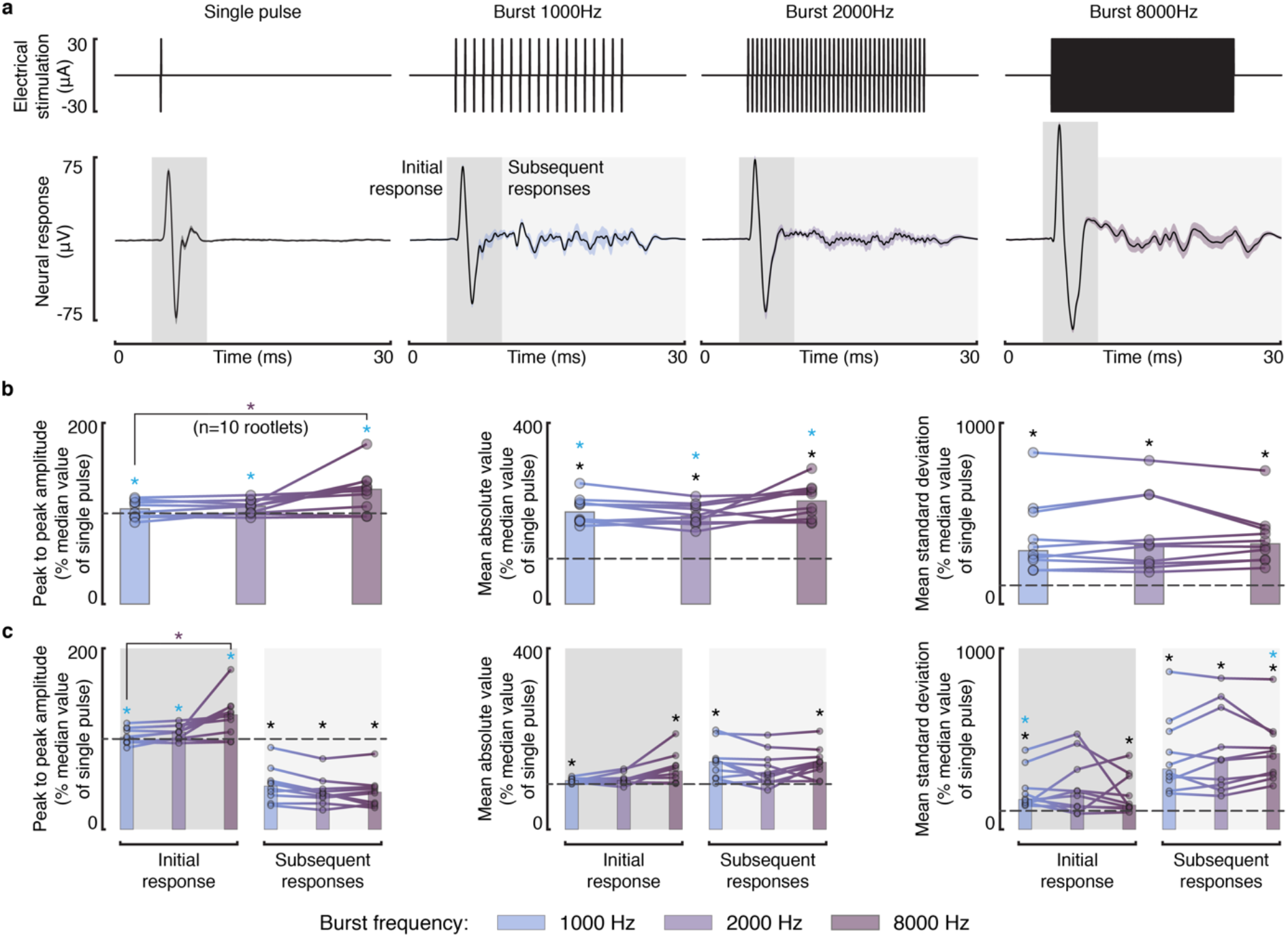
Effect of constant-amplitude high-frequency bursts on heterogeneous populations of rat afferents. **a**, Average (± standard deviation) of the neural responses induced by single pulses of stimulation (as in Figure3c), and by constant-amplitude high-frequency bursts delivered at 1000, 2000, and 8000 Hz, respectively from left to right, in a representative extracted rat dorsal rootlet (burst duration: 20 ms; pulse width: 50 μs). Dark-gray shaded areas highlight the initial neural response (−1 to 5 ms from the stimulation onset); light-gray shaded areas highlight subsequent responses (5 to 25 ms from the stimulation onset). **b**, Scatterplots reporting, from left to right, the median of the peak to peak amplitude, mean absolute value, and mean standard deviation of the neural responses induced by the tested burst conditions, for all (n=10) the tested rootlets. For each rootlet, statistics are normalized with respect to the median value of single pulse stimulation (highlighted by the dashed lines). Lines connecting different burst conditions indicate the results for different rootlets. Bars represent the median of each statistic across the tested rootlets. Black stars indicate a significant difference between the starred burst condition and 1 (i.e., the median value of single pulse stimulation); p<0.05, two-sided Wilcoxon signed rank test with Bonferroni correction for multiple comparisons. Cyan stars indicate a significant difference between the starred burst condition and BioS bursts delivered at the same frequency (Figure 3e); violet asterisks indicate a significant difference between two burst conditions; p<0.05, two-sided, Wilcoxon signed rank test with Bonferroni correction for multiple comparisons. **c**, Scatterplots reporting the statistics of the initial neural response and of the subsequent responses independently. Instead of the mean absolute value, we reported the sum of absolute values to avoid accounting for the difference length of these two responses. Stars and bars follow the same conventions as in **b**.

As opposed to BioS, constant-amplitude bursts induced a strong, synchronous response during the first few pulses of stimulation, which resembled the compound action potential induced by a single pulse of stimulation (Figure 4a). Indeed, we did not observe any significant differences between the peak-to-peak amplitudes of the responses induced by a constant amplitude burst and single pulse stimulation (Figure 4b, * p<0.05, two-sided Wilcoxon signed rank test with Bonferroni correction for multiple comparison). Interestingly, bursts at 8000 Hz induced responses with higher peak-to-peak amplitudes compared with those induced at 2000 Hz. This is in agreement with our model’s prediction, suggesting that increasing the stimulation frequency above a certain threshold recruits a larger number of fibers due to summation effects.

The strong, synchronous response induced at the beginning of each constant amplitude burst was followed by multiple weak, asynchronous responses generated throughout the entire burst duration (Figure 4a and c). Consequently, the mean absolute value and the mean standard deviation of the induced responses were significantly higher than those obtained with single pulse stimulation (Figure 4b). In particular, these additional responses, taken together, were of similar magnitude to the initial compound action potential, and matched the activity induced by BioS bursts, suggesting that constant-amplitude bursts elicited significant additional action potentials in the recruited fibers after the initial response (Figure 4c). The peak-to-peak amplitude of these subsequent responses was significantly lower than those induced by a single stimulation pulse, but was not different from those induced by BioS, indicating that these additional responses were similarly asynchronous (Figure 4c, * p<0.05, two-sided Wilcoxon signed rank test with Bonferroni correction for multiple comparison).

These results suggest that constant-amplitude high-frequency bursts induce responses that are only partially desynchronized, confirming our hypothesis that modulating the stimulation amplitude of each burst is essential for inducing an asynchronous, stochastic, and controllable neural response.

### BioS burst maximum amplitude controls the number of recruited fibers

We explored whether modulating the maximum amplitude reached within a BioS burst would allow precise control over the number of recruited fibers, the same way modulating pulse amplitude does in the case of pulsed stimulation. We recorded the neural responses of 3 dorsal rootlets to BioS bursts and single pulses delivered at stimulation amplitudes ranging from 120% to 200% of the threshold amplitude required to induce a neural response (Figure 5a). The first pulse within a BioS burst was also set to the threshold amplitude.

**Figure 5.**
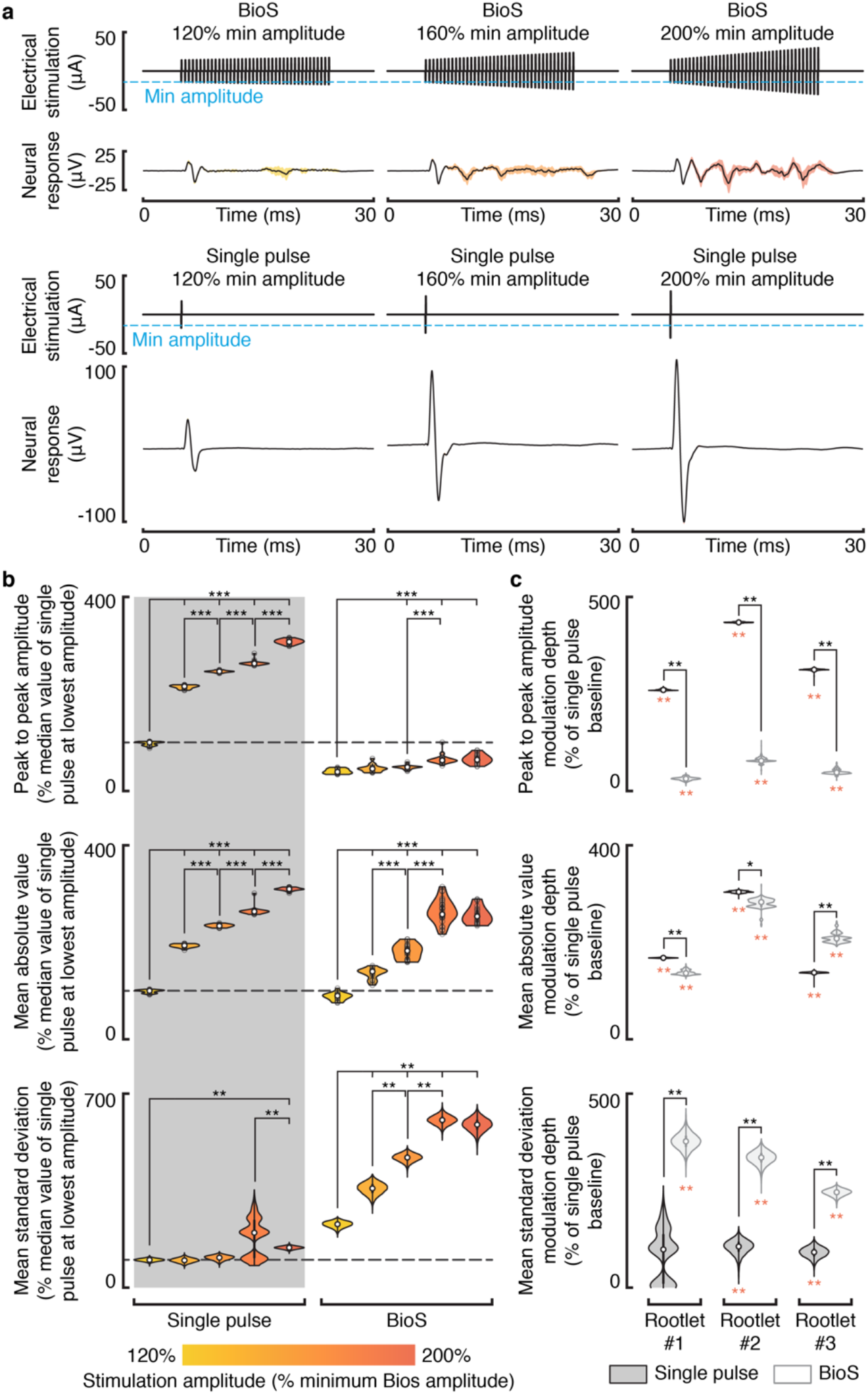
BioS burst amplitude modulation. **a**, Average (± standard deviation) of the neural responses induced by BioS bursts (top) and single pulses of stimulation (bottom) for 3 representative amplitudes, in rootlet #1 (burst duration: 20 ms; burst frequency: 2000 Hz; pulse width: 50 μs). Stimulation amplitudes are reported in % of the minimum amplitude used within a BioS burst (as highlighted by the cyan dashed line), which we set to the lowest amplitude necessary to induce a neural response. **b**, Violin plots reporting, from top to bottom, the peak to peak amplitude, mean absolute value, and mean standard deviation of the neural responses induced by the tested stimulation conditions, for rootlet #1 (n = 115 responses). Each statistic is normalized with respect to the median value of single pulse stimulation, as highlighted by the dashed lines. The violin plots for the peak to peak amplitude and the mean absolute value report the sampled distributions. Violin plots for the mean standard deviation report the bootstrapped distribution (n iterations = 10,000). Stars indicate a significant difference: ***, p<0.0001; **, p<0.005; two-sided Wilcoxon rank sum test with Bonferroni correction for multiple comparisons. The same plots for rootlets #2 and #3 are reported in Supplementary Figure 3. **c**, Violin plots reporting the bootstrapped (n iterations = 10,000) modulation depth of each statistic (computed as the difference between the maximum and the minimum of the medians calculated over the tested stimulation amplitudes), for all the tested rootlets. Values are normalized with respect to the baseline of single pulse stimulation, i.e., the median value of the lowest tested amplitude. Dark grey plots report the statistics for single pulse stimulation, light grey plots for BioS. Orange stars (below the violin plots) indicate a significant modulation (modulation > 0); **, p<0.005; one-sided bootstrap test with Bonferroni correction for multiple comparisons. Black stars indicate a significant difference between single pulse and BioS; **, p<0.005; *, p<0.05; two-sided, bootstrap test with Bonferroni correction for multiple comparisons.

We found that the mean absolute value of the responses to both BioS bursts and single pulses progressively increased with increasing stimulation amplitude (Figure 5b and Supplementary Figure 3). We found similar, yet significantly different, modulation depths (computed as the difference between the maximum and the minimum of the medians calculated over the tested stimulation amplitudes) for the two stimulation conditions (Figure 5c, across rootlets average depth of 261% and 254% for BioS and single pulse stimulation, respectively). In the case of single pulses, we observed an increase in peak to peak amplitude in response to higher stimulation amplitudes (median increase > 260.3% across all the tested rootlets), consistent with the fact that an increasing numbers of fibers were being recruited synchronously (Figure 5b, c and Supplementary Figure 3). On the other hand, BioS bursts demonstrated only modest increases in peak to peak amplitude when using higher stimulation amplitudes (median increase < 78.4% across all the tested rootlets). Finally, in the case of BioS bursts, the mean standard deviation of the response robustly increased with higher stimulation amplitudes (median increase > 245.1% across all the tested rootlets), while only modestly increasing in the case of single pulse stimulation (median increase < 106% across all the tested roots, statistically significant increase in only 2 rootlets, p<0.005).

These results demonstrate that BioS bursts can be used to recruit a controllable number of fibers, while maintaining strong desynchronization, and that the higher the number of recruited fibers, the larger the variability of the induced response.

### Tradeoff between synchronicity and firing stability during continuous stimulation

Given the limited ability of axons to keep up with high frequency stimulation profiles^27–30^, we reasoned that repeatedly delivering BioS bursts (continuous stimulation) might lead to rapid depression of the neural responses. Based on this assumption, we further hypothesized that interleaving consecutive bursts with a period of no-stimulation (duty cycling) might be necessary to overcome this limitation. To test these hypotheses and evaluate whether BioS can induce sustained asynchronous neural activity, we stimulated 3 dorsal rootlets with 10-seconds long, trains of BioS bursts with duty cycles of 50%, 70% and 90% and burst repetition frequency of 20 Hz (Figure 6a). For comparison, we also stimulated the extracted rootlets with pulsed stimulation delivered at 20 Hz.

**Figure 6.**
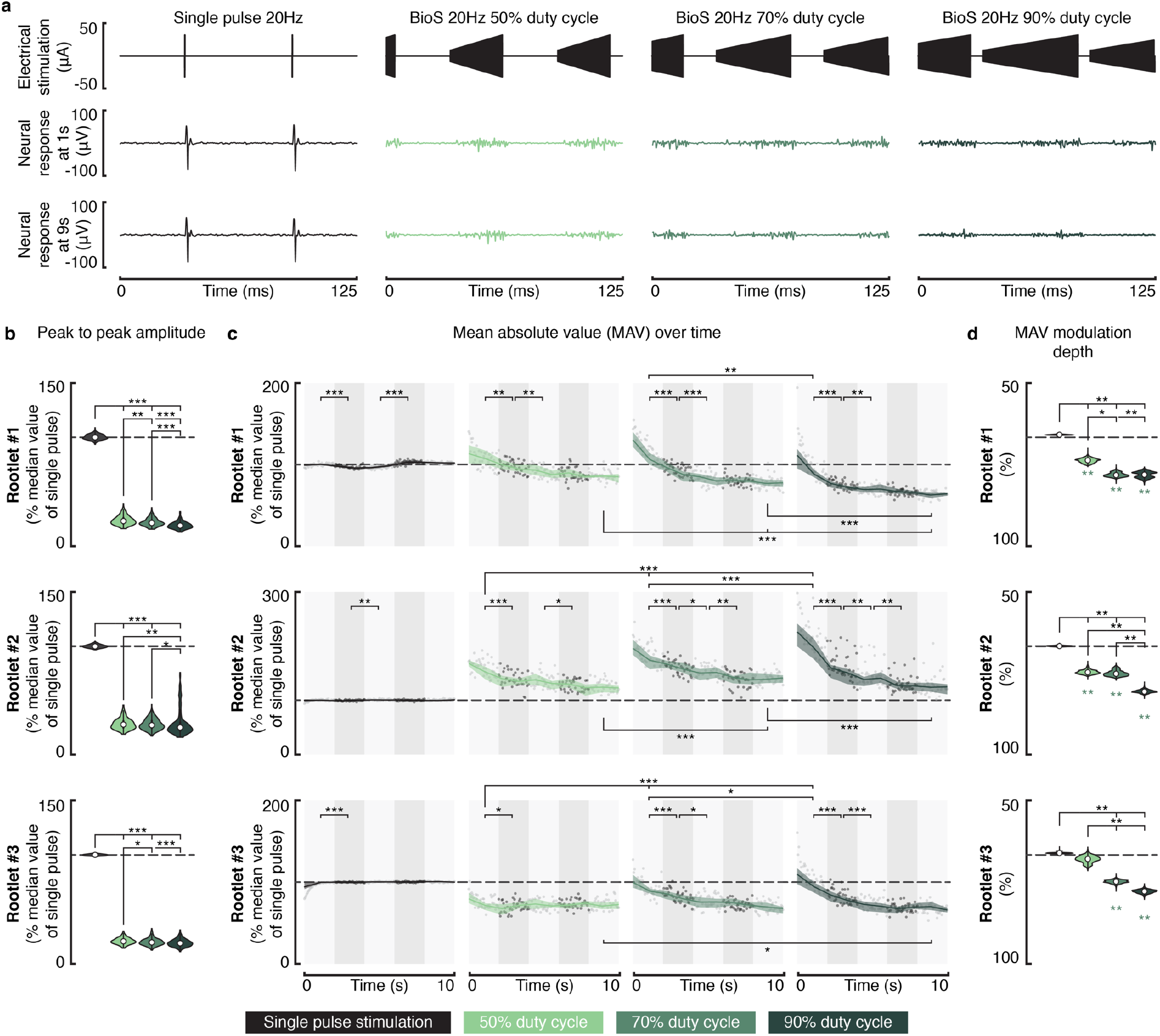
Temporal stability of the neural activity induced by continuous BioS profiles. **a**, Representative neural responses induced by 20 Hz pulsed stimulation (left), and by 20 Hz BioS profiles with different duty cycles (50%, 70%, and 90%, respectively from left to right), at 1 s (top) and 9 s (bottom) after the stimulation onset, in rootlet #1. Each stimulation profile was delivered for a duration of 10 s. **b**, Violin plots reporting the peak to peak amplitude of the neural responses induced by each stimulation condition, for all the tested rootlets (n = 800 responses, for each rootlet). Statistics are normalized with respect to the median value of single pulse stimulation, as highlighted by the dashed lines. **c**, Temporal analysis of the induced responses’ mean absolute value. Scatterplots report the time evolution of this statistic. Solid lines and shaded areas represent the median values and the IQR, respectively. Shaded grey rectangles indicate the 2-s intervals on which the statistical analysis is performed. **b**, **c**, Stars indicate a significant difference: ***, p<0.0001; **, p<0.005; *, p<0.05; two-sided Wilcoxon rank sum test with Bonferroni correction for multiple comparisons. **d**, Violin plots report the bootstrapped (n iterations = 10,000) modulation of the mean absolute value, computed as the difference between the median values obtained from the last 2 and the first 2 seconds of recording. Statistics are normalized with respect to the median of the mean absolute value computed during the first 2 s of recording. Green stars indicate a significant negative modulation (modulation < 0); p<0.005; one-sided bootstrap test with Bonferroni correction for multiple comparisons. Black stars indicate a significant difference between the tested stimulation conditions; **, p<0.005; *, p<0.05; two-sided, bootstrap test with Bonferroni correction for multiple comparisons.

All the tested BioS conditions induced neural responses which were significantly less synchronized than pulsed stimulation (lower peak to peak amplitude, p<0.0001, two-sided Wilcoxon rank sum test with Bonferroni correction for multiple comparisons, Figure 6b). Using a duty cycle of 50% consistently induced the responses with the highest peak to peak amplitude. This result is consistent with the hypothesis that BioS bursts distribute neural recruitment over their entire duration thereby evoking more synchronized responses when their duration is shortened. This also suggests that neural synchronicity can be modulated by changing the burst duration.

Regarding the mean absolute value, we observed a decrease over time during BioS, which was absent during pulsed stimulation (Figure 6c and d). For all the tested duty cycles and rootlets, this decrease reached a plateau after 8 seconds at most (Figure 6c, no significant decrease between samples collected 6-8 seconds after the stimulation onset and those collected 8-10 seconds after). Stimulation profiles with higher duty cycles generally produced stronger responses at the beginning of the trial compared with lower duty cycle profiles (Figure 6c), but also suffered from a stronger depression of the responses (Figure 6d). When BioS was delivered with a duty cycle of 50%, the maximum reduction of the response over the 3 tested rootlets was of 24% (bootstrapped median, n iterations = 10,000), while this increased to 35% and 42% when the duty cycle was incremented to 70 and 90%, respectively (Figure 6d). This finding not only suggests that high duty cycle BioS profiles induce stronger neural fatigue, but also that longer bursts initially evoke a greater number of action potentials in the recruited population of fibers.

Taken together these results show that BioS can induce sustained asynchronous neural activity, while also highlighting a tradeoff between the level of desynchronization that can be achieved and the temporal stability of the evoked responses. Specifically, while high duty cycle BioS bursts will cause the most desynchronization (since the bursts are longer), they will also more strongly attenuate the neural response over time.

### BioS repetition frequency regulates the firing rate of the recruited fibers

Finally, we explored whether modulating the BioS repetition frequency would enable control over the firing rate of the recruited fibers, analogously to the case of pulsed stimulation. For this, we tested the effect of BioS and pulsed stimulation at frequencies ranging between 10 and 80 Hz (n = 3 rootlets). For all the tested frequencies, we delivered BioS with a 70% duty cycle and recorded the responses for 10 s. Figure 7a reports illustrative neural responses of rootlet number 1 to both BioS and pulse-based stimulation at 20, 40, and 80 Hz.

**Figure 7.**
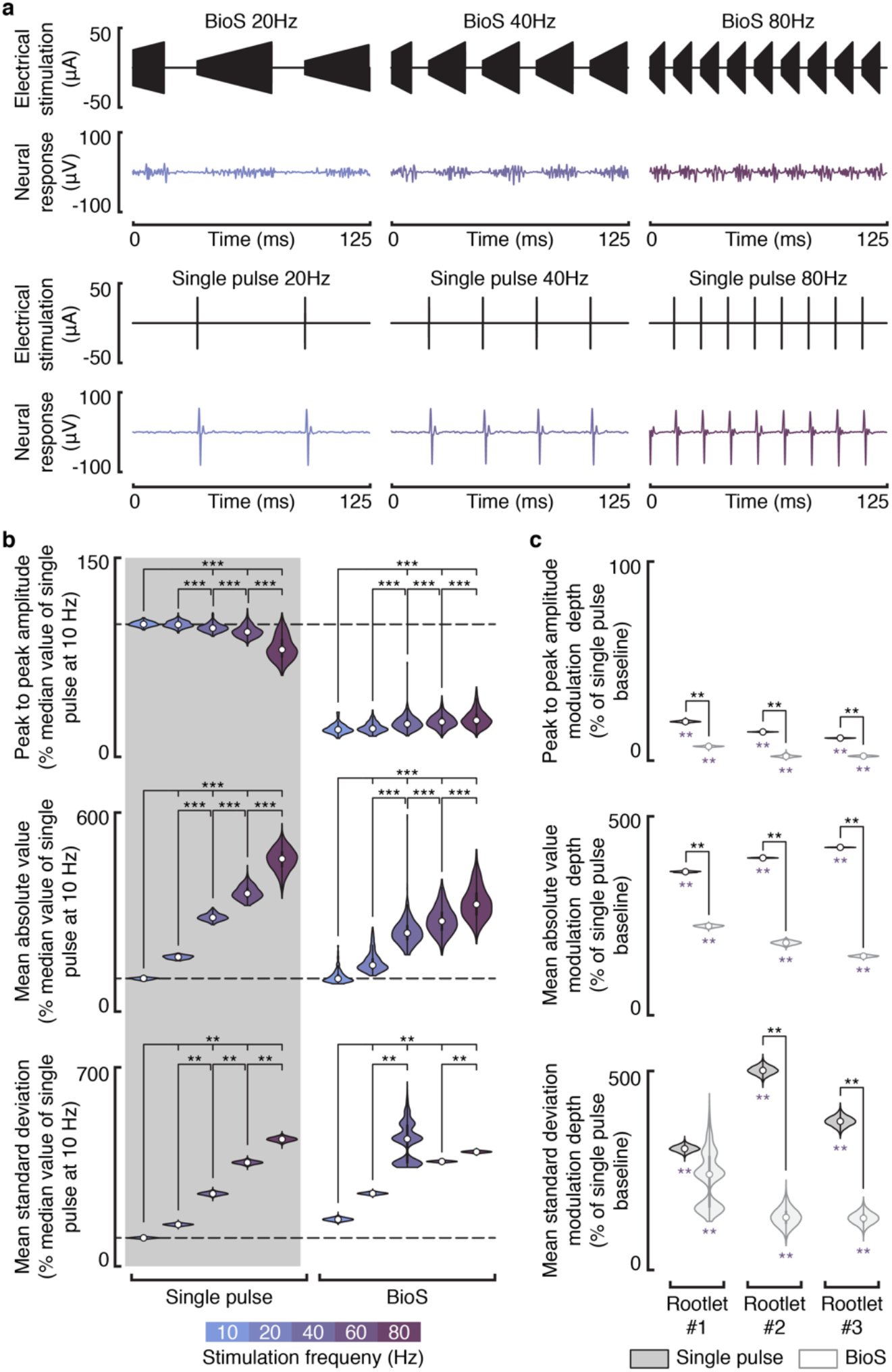
BioS repetition frequency modulation. **a**, Representative neural responses induced by BioS (top) and pulsed stimulation (bottom) delivered at 20, 40, and 80 Hz (from left to right), in rootlet #1 (burst frequency: 2000 Hz; burst duty cycle: 70%; pulse width: 50 μs). **b**, Violin plots reporting, from top to bottom, the peak to peak amplitude, mean absolute value, and mean standard deviation of the neural responses induced by the tested stimulation conditions, for rootlet #1 (n = 37,713 responses). Each statistic is normalized with respect to the median value of single pulse stimulation, as highlighted by the dashed lines. The violin plots for the peak to peak amplitude and the mean absolute value report the sampled distributions. Violin plots for the mean standard deviation report the bootstrapped distribution (n iterations = 10,000). Stars indicate a significant difference: ***, p<0.0001; **, p<0.005; two-sided Wilcoxon rank sum test with Bonferroni correction for multiple comparisons. The same plots for rootlets #2 and #3 are reported in Supplementary Figure 4. **c**, Violin plots reporting the bootstrapped (n iterations = 10,000) modulation depth of each statistic (computed as the difference between the maximum and the minimum of the medians calculated over the tested stimulation frequencies), for all the tested rootlets. Values are normalized with respect to the baseline of single pulse stimulation, i.e., the median value of the lowest tested frequency. Dark grey plots report the statistics for single pulse stimulation, light grey plots for BioS. Violet stars (below the violin plots) indicate a significant modulation (modulation > 0); **, p<0.005; one-sided bootstrap test with Bonferroni correction for multiple comparisons. Black stars indicate a significant difference between single pulse and BioS; **, p<0.005; *, p<0.05; two-sided, bootstrap test with Bonferroni correction for multiple comparisons.

Increments in stimulation frequency induced a progressive increase in the mean absolute value of the responses evoked by both BioS and pulsed stimulation, in all the tested rootlets (Figure 7b and Supplementary Figure 4, *, p**<**0.05, **, p<0.005, ***, p<0.0001, two-sided Wilcoxon rank sum test with Bonferroni correction for multiple comparisons). This modulation, however, was significantly larger in the case of pulsed stimulation, reaching a maximum modulation depth of 422%, as opposed to 224% in the case of BioS (Figure 7c, **, p<0.005, bootstrap test with Bonferroni correction for multiple comparisons). Increasing the frequency also led to a steady, yet weak, decrease in the peak-to-peak amplitude of the responses generated by pulsed stimulation, but not by BioS (Figure 7b, c and Supplementary Figure 4). Finally, higher stimulation frequencies led to responses with progressively greater standard deviation (Figure 7b and Supplementary Figure 4).

Overall, these results confirm that adjusting the repetition frequency of continuous BioS profiles enables control over the firing rate of the recruited fibers. Furthermore, these results also highlight differences between BioS and pulsed stimulation in the level of modulation that can be achieved.

## Discussion

We developed a biomimetic electrical stimulation strategy (BioS) that induces controllable, strongly asynchronous, and stochastic neural activity, thus overcoming the limitations of pulsed stimulation. Instead of single pulses, the proposed strategy uses high-frequency amplitude-modulated bursts as fundamental stimulation units (Figure 2a). Two hypotheses underpinned this design. The first was that ramping up stimulation amplitude within a burst would ensure the progressive and desynchronized recruitment of target fibers. Initially, low amplitude pulses would recruit large diameter fibers and fibers very close to the stimulating electrode. Then, as the amplitude of subsequent pulses increases, smaller diameter fibers or fibers more distant from the electrode would be recruited as well. Finally, the last few pulses in the burst, with the highest current amplitudes, would activate the smallest and most distant fibers in the target population. The second hypothesis was that the limited ability of neurons to follow high-frequency stimulation profiles^27–30^ would prevent fibers from firing within each burst. Therefore, a BioS burst would induce a similar number of action potentials as a single stimulation pulse, while spreading recruitment throughout its duration. Here we discuss the significance of our results, some key BioS parameters and how to tune them, and the implications of this novel stimulation strategy for the field of neuromodulation.

### BioS induces asynchronous, stochastic, yet controllable, neural activity

We leveraged a computational model and ex vivo experiments to validate the hypotheses supporting the proposed stimulation strategy. We obtained compelling evidence suggesting that BioS bursts induce asynchronous neural activity by distributing fiber recruitment throughout their duration, and found BioS configurations that led to similar overall levels of neural activity compared to pulsed stimulation. These findings corroborated our fundamental hypotheses.

Both computational modeling and ex vivo experiments demonstrated that BioS induces stochastic neural activity. We propose that this phenomenon arises from the presence of high-frequency pulses within a BioS burst. Indeed, when using burst frequencies above approximately 2kHz, neuronal membranes have insufficient time to return to their resting potential between consecutive pulses in a burst. This implies that each stimulation pulse induces a membrane depolarization that affects the fiber’s response to the next pulse (as demonstrated in Figures 2b and 4). Consequently, the exact moment a fiber is recruited depends on its membrane potential at the beginning of a burst, which can vary slightly from burst to burst due to natural membrane potential fluctuations, causing the response of an individual fiber to BioS bursts to be highly variable.

We showed that both the number of recruited fibers and the induced firing rate can be controlled by modulating BioS amplitude and repetition frequency, analogously to the case of pulsed stimulation. Interestingly, although increasing the stimulation amplitude used for BioS bursts resulted in progressive increases in average neural activity, this was also accompanied by small increases in peak to peak amplitude (Figure 5b, c and Supplementary Figure 3). This was likely caused by the fact that when burst amplitude increases, the modulation range likewise increases. For instance, at 120% of the threshold amplitude, a BioS burst is modulated from threshold to 20% above threshold in a fixed amount of time. If the amplitude is increased to 180% of the threshold, the burst is modulated from threshold to 80% above threshold in the same amount of time. Since the number of individual pulses within a BioS burst only depends on the intra-burst frequency (fixed at 2 kHz in the ex vivo experiments) and the burst duration, it is unaffected by amplitude modulation. Consequently, the difference in amplitude between consecutive pulses within a burst becomes larger as the modulation range increases, which in turn makes it more likely for two or more fibers to be recruited by the same pulse. This is consistent with the small increase in peak to peak amplitude observed at larger BioS amplitudes.

During frequency modulation, we observed that BioS induced progressively higher average neural activity, confirming that adjusting the BioS repetition frequency allows fine-tuning of the recruited fibers’ firing rate (Figure 7). However, the magnitude of this modulation was significantly lower compared with pulsed stimulation. We propose two concurrent mechanisms to explain this observation. First, as seen in Figure 6, shorter bursts induce weaker neural activity compared to longer bursts (lower mean absolute value). Therefore, since an increased stimulation frequency is associated with a shortened burst duration, a lower number of action potentials will be generated during a burst repeated at high frequencies. This is also consistent with the nearly absent modulation in the responses’ peak-to-peak amplitude observed when increasing the BioS repetition frequency. Indeed, if the same number of action potentials were induced in a shorter period of time (i.e., when high BioS repetition frequencies are used), a higher peak-to-peak amplitude would be expected, as discussed in the case of amplitude modulation. However, since this was only the case in one of the three tested rootlets (Figure 7b and Supplementary Figure 4), it is likely that shorter bursts induced weaker overall neural activity. We believe that this phenomenon is caused by a lower number of action potentials in the recruited fibers, rather than a reduced number of recruited fibers. Second, the mean absolute value likely underestimates the overall neural activity when multiple action potentials are asynchronously induced in a brief period of time. Indeed, when using short burst durations, the probability that the electrical signals originating from two action potentials occurring in quick succession will interfere with each other increases (e.g. depolarization and hyperpolarization of two fibers cancelling each other out at the recording site). As the probability of such interferences increases, the magnitude to which the mean absolute value of the measured potential underestimates the overall induced neural activity likewise increases. Consequently, we hypothesize that the observed mean absolute value modulation during BioS is underestimated, and that this effect is strongest for very high repetition frequencies.

Interestingly, increasing single pulse stimulation frequency led to a weak, but significant decrease in the responses’ peak-to-peak amplitude. This result likely stems from the refractory period of the target fibers. Indeed, when increasing the stimulation frequency above a certain threshold, it is likely that the period between two consecutive stimulation pulses becomes shorter than the refractory period of some of the fibers in the population, thus reducing the probability of all fibers responding to each stimulation pulse. This interpretation is consistent with the observed increase in mean standard deviation at higher frequencies, which suggest that consecutive pulses recruit a different number of fibers. These results also point out that stimulating with simple high frequency trains of pulses — at frequencies above the maximum firing rate of the target fibers, but below the kHz range to avoid the types of blocking effects exploited by BioS (e.g., at 300 Hz) — should also lead to asynchronous neural activity. Indeed, since fibers will not be able to fire at each stimulation pulse, they will slowly desynchronize due to their slightly different refractory periods. However, this strategy would elicit uncontrollable neural firings, since the population firing rate would be tied to the maximum firing rate of the fibers, making this approach unviable for most applications.

These results demonstrate that BioS can induce asynchronous, stochastic, yet controllable, neural activity. This level of control over the recruited fibers is essential for generating biomimetic patterns of neural activity, and cannot be achieved with pulsed stimulation.

### Key BioS parameters and design features

Some BioS conditions induced significantly higher levels of neural activity compared to pulsed stimulation (Figure 3d-e), suggesting that certain BioS parameters caused some fibers to respond more than once per burst. Our modeling results confirmed that fibers could fire more than once in response to a single BioS burst, and suggested that these events depend on the burst frequency, with repeated responses almost exclusively observed when using burst frequencies below 2000 Hz (Figure 2e). These modeling results were not fully corroborated during ex vivo experiments, where bursts frequencies as low as 1000 Hz rarely induced consecutive firings in the same fibers. We hypothesize that this discrepancy was the result of the different temperatures used in both experiments, with the simulations performed at 37 degrees, and the ex vivo experiments performed at room temperature (approximately 23 degrees). Adjusting the temperature in the computational model brought simulations in line with the experimental results (Supplementary Figure 5). Overall these results highlight the importance of properly adjusting the BioS burst frequency to minimize the number of times a fiber is recruited within a burst, which is essential to enable precise control over the induced neural activity. Nevertheless, it is important to note that even if a fiber is recruited multiple times within a burst, its firing rate would remain controllable (by adjusting the BioS repetition frequency) as long as it remains lower than the maximum firing rate allowed by the fiber’s refractory period.

We demonstrated that amplitude-modulation of the high frequency carrier is essential for inducing asynchronous, controllable neural activity. In particular, we found that constant-amplitude high-frequency bursts induced neural activity consisting of both a synchronous response, of equal or greater magnitude to that induced by a single stimulation pulse delivered at the same amplitude, followed by several asynchronous responses. These results suggest that constant-amplitude bursts induce responses that are more synchronous and involve a higher number of action potentials in the recruited fibers, compared with BioS bursts. This finding is largely consistent with previous high-frequency studies, reporting similar transient neural activations when constant-amplitude high-frequency stimulation profiles were used for nerve conduction block^28,34^, but minimal activations when the stimulation amplitude was gradually ramped-up^30^.

Although we only tested BioS bursts with linearly increasing amplitude, more complex amplitude-modulation profiles might further desynchronize the induced neural activity. To maximize neural desynchronization, each pulse within a burst should recruit the same number of fibers, so that fiber recruitment is evenly distributed throughout the entire burst duration. As can be seen in Figures 2d and 3, a linear increase in stimulation amplitude leads to inhomogeneous fiber recruitment, with a distribution skewed towards the end of the burst. The shape of the fiber recruitment function depends on several factors, including the conductive properties of the stimulated tissue, the spatial arrangement of the targeted fiber population, and the distribution of fiber diameters. Assuming a homogenous space populated by equally distributed, identical fibers, and a point source electrode, a linear increase in stimulation amplitude during a BioS burst would lead to a cubic fiber recruitment function, i.e., consecutive stimulation pulses within a burst would recruit a cubically-increasing number of fibers (this phenomenon is simply caused by the spherical propagation of the electric field around the electrode). Therefore, to flatten this recruiting function and maximize neural desynchronization, the amplitude-modulation profile should follow a cubic root function. In realistic scenarios, the optimal amplitude-modulation profile should be computed as a function of the target tissue and electrode properties.

In addition to the stimulation amplitude modulation function, fine tuning the range of stimulation currents is also essential to ensure optimal neural desynchronization. Using a minimum amplitude lower than the smallest fiber threshold would reduce the period of time during which fiber recruitment can be desynchronized, as only a portion of the pulses in a burst would effectively recruit any fibers. A similar effect would occur if the maximum amplitude were larger than the highest fiber threshold, whereby a certain number of the highest amplitude pulses within the burst would not recruit any further fibers. Inversely, if a high minimum amplitude is used, the first pulse within a burst would recruit multiple fibers and lead to a synchronized response. Therefore, ideally the minimum stimulation amplitude within a BioS burst should be calibrated to only recruit the neural structures with the lowest thresholds, while the maximum amplitude should be set at the lowest amplitude necessary to recruit the entire target population.

Finally, we showed that BioS can be used to induce continuous asynchronous neural activity, but that the stability of the induced response depends on the stimulation duty cycle. In particular, a high duty cycle might lead to strong depression of the induced responses. We also found that adjusting the BioS duty cycle allows control over the level of desynchronization, with high duty cycle stimulation inducing more asynchronous responses compared with low duty cycle stimulation.

### Biomimetic stimulation of the nervous system

Electrical stimulation of the nervous system is a valuable tool, allowing researchers to deliver information to the nervous system. However, as highlighted in this work, current strategies fail to generate signals that “speak” the language of the nervous system: the neural activity induced by pulsed electrical stimulation is highly repeatable and synchronous, while naturally occurring activity is stochastic and highly asynchronous. BioS has the potential to reduce this nonconformity, and thus improve our ability to communicate with the nervous system. In this section, we explore the impact of the proposed stimulation strategy on selected neurostimulation applications.

Functional electrical stimulation (FES) uses pulsed stimulation to control muscle activity. However, because neural responses induced by pulsed stimulation are strongly synchronous, FES induces jerky movements and higher muscle fatigue, compared with physiological recruitments^24,25^. During sustained physiological muscle contractions, motoneurons fire asynchronously at a rate of 6 to 8 impulses per second^9^. At these frequencies, asynchronous firing is essential to produce the smooth force profiles underlying natural movements, by ensuring the generation of a continuous force. Consequently, delivering FES with pulsed stimulation at 6 Hz would lead to uncontrollable spasmodic movements due to the synchronous contraction of motor units. To compensate for this, current FES paradigms use much higher frequencies (between 20 and 40 Hz). This enables the generation of sustained muscle contractions, but at the expenses of increased muscle fatigue^9,24,25^. Using BioS during FES would thus allow a decrease in muscle fatigue and much finer control over the induced muscle contractions, potentially increasing the benefits provided by this technique to patients with motor disabilities.

Another application in which BioS could be extremely helpful is sensory feedback restoration^20^. Indeed, electrical stimulation of cutaneous afferents typically induces unnatural sensations. In fact, evoked sensations are often reported as vibration or fluttering, which would be consistent with the synchronicity of the restored sensory signals. Two recent studies demonstrated the importance of delivering biomimetic patterns of electrical stimulation to restore the sense of touch in trans-radial amputees during control of a myoelectric hand prosthesis^8,17^. Using peripheral nerve stimulation, these studies showed that modulating the frequency and amplitude of pulsed stimulation over time to mimic the natural firing patterns of touch induces more natural sensations. However, the ability to induce highly biomimetic patterns of neural activity was likely hampered by the non-physiological, synchronous nature of the elicited neural responses. Indeed, even if the population firing rate closely matched naturally occurring firing patterns, the induced activity remained time-locked to each stimulation pulse. Therefore, it is possible that using BioS to further improve the level of biomimicry of the induced neural activity would evoke even more natural sensations.

In this work, we focused on nerve fibers. Nevertheless, BioS would likely elicit the same effects in any type of neuron. Indeed, the mechanisms underlying the proposed strategy only rely on fundamental membrane properties, which are common to all neural structures. Thus, applications involving cortical or subcortical stimulation may also benefit from BioS^35–38^. However, it is important to note that some applications likely benefit from synchronicity. For instance, during spinal cord stimulation for gait restoration, the efficacy of certain electrical stimulation protocols depends on the magnitude of the post-synaptic potentials induced in targeted motoneurons by the recruitment of proprioceptive fibers^39^. It is to be expected that higher synchronicity in the afferent activity would cause larger post-synaptic changes, making BioS a poor candidate for this specific application.

In conclusion, BioS is an addition to the neurostimulation toolkit which enables researchers to control a new parameter when recruiting neural structures: synchronicity. Inducing synchronous neural responses will no longer be the default choice. Instead, researchers will be able to choose the strategy that best fits their needs and application, thus opening up interesting new opportunities.

## Materials and methods

### Computational model of a bundle of nerve fibers during electrical stimulation

The simulated bundle was composed of a heterogeneous population of myelinated fibers. These were modeled as simple myelinated axons with no cell body, and had two components: a biophysical compartmental model, which we used to evaluate the effect of the stimulation on the membrane potential; and a propagation model, which we used to simulate action potential propagation.

The biophysical model was implemented using the NEURON simulation environment^40^, and was based on the multicompartment cable model proposed by McIntyre and colleagues^41^. This model was shown to accurately predict the excitation properties of mammalian myelinated fibers to both pulsed^41^ and high-frequency^29^ extracellular electrical stimulation. Ten compartments were used to model the attachment (MYSA), paranode (FLUT), and internode (STIN) fiber segments between successive nodes of Ranvier (Figure 1a). Segment dimensions are based on realistic morphological measurements, which were obtained from 9 fibers with diameters ranging from 5.7 to 16.7 µm^41^. To generate additional fibers beyond those that were observed, we used a quadratic polynomial fit over the sampled dimensions. Independently of the diameter, fibers were modeled with 41 nodes of Ranvier (441 segments in total). Thus, fibers with different diameters also had different lengths.

To simulate action potential propagation along a given fiber length, we implemented a propagation model. When an action potential was induced in the biophysical model, its propagation velocity was computed and used to estimate the time the same action potential would take to travel along the imposed fiber length. In this way, even if the biophysical model was only a few centimeters long (which is enough to evaluate the effect of electrical stimulation on the membrane), we were able to estimate action potential propagation along arbitrarily long fibers, without drastically increasing the computation time.

We used this fiber model to create a heterogeneous population of fibers with properties tuned to match the scenario of upper-limb peripheral nerve stimulation. Specifically, we created a population of 100 group-II afferents, with diameters following a normal distribution with mean of 9.2 µm and standard deviation of 0.5 µm, and length of 40 cm. Fibers were distributed uniformly in a circle of radius 100 µm, with a random longitudinal shift to avoid lining up all nodes of Ranvier.

To evaluate the effect of electrical stimulation, we considered a point source electrode positioned in the center of the modeled population, at 7.25 mm from the beginning of the fiber, and an infinite, homogeneous, isotropic medium. We then simulated an electrical stimulation waveform and computed the external potential at each model compartment, using the formula 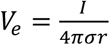, where *I* denotes the injected current, σ the conductivity of the medium (σ = 2 S/m), and *r* the distance from the electrode to the considered compartment.

When evaluating the effect of pulsed stimulation (frequency: 40 Hz; amplitude: 0.8 mA; pulse width: 50µs; Figure 1c-e) and BioS (repeating frequency: 40 Hz; burst frequency: 3500 Hz; amplitude: 0.8 mA; pulse width: 50µs; Figure 2c-e), we ran simulations lasting 500 ms, in which the stimulation was turned on only 15ms after the start of the simulation. This ensured that all membrane potentials could reach their resting values. To reduce the computation time, simulations performed to characterize the effect of different BioS burst frequencies (1000–8000 Hz) and amplitudes (0.1–0.9 mA, Figure 2b) were performed over 200 ms. Simulations in Figure 1 and 2 were performed at body temperature (37 °C); simulations in Supplementary Figure 5, were performed at room temperature (23 °C).

### Computational model of a bundle of nerve fibers exhibiting natural neural activity

Natural neural activity — which we used to evaluate the ability of a stimulation waveform to produce biomimetic firings — was modeled using homogeneous Poisson processes. For each modeled fiber, the interval between consecutive action potentials followed a negative exponential distribution with a fixed mean. This widely used approach provides a convenient approximation of the stochastic properties of natural neural firings^42^. When comparing electrically-induced and natural firings, we simulated natural neural activity with a mean firing rate matching the one induced by the tested electrical stimulation condition (pulsed stimulation in Figure 1c-e, and BioS in Figure 2c-e).

### Analysis of the simulated neural activity

The population neural activity was estimated by convolving the induced action potentials — recorded at the end of the fibers and expressed as trains of Dirac delta functions — with a Gaussian wavelet (duration: 1 ms; standard deviation: 0.33 ms), and by subsequently adding up the results (Figures 1c and 2c). The neural activity was then binned in windows of 25 ms (Figure 1d and 2d), starting 8.5 ms after the stimulation onset (estimated minimum period of time required for an action potential induced below the electrode to propagate along the whole length of a simulated fiber). Three statistics were computed over the extracted responses: the mean absolute value, the peak to peak amplitude, and the mean standard deviation. The mean absolute value measured the magnitude of the response, i.e., the number of action potentials generated in the entire population (greater values in this statistic indicates higher levels of neural activity). The peak to peak amplitude measured the response synchronicity; assuming that a similar number of action potentials is induced, the summation of multiple synchronous responses would cause higher maximum values. Finally, the mean standard deviation measured the response variability. In addition to these three statistics, we also extracted the average number of times a recruited fiber fired and the overall number of recruited fibers, to assess fiber recruitment.

### Rat dorsal rootlets extraction

All procedures were approved by the Veterinary Office of the canton of Geneva in Switzerland, and were in compliance with all relevant ethical regulations. Dorsal rootlets were extracted from 4 adult male Lewis rats, following the procedures described in ^33^. Briefly, rat spinal cords were exposed and nerve roots were cut at the exit foramen. These were then dissected under a microscope to obtain a nerve strand, which was further teased into rootlets approximately 60 µm in diameter (Supplementary Figure 1). A total of 16 dorsal rootlets were extracted: 10 for the experiments reported in Figure 3 and 4, 3 for the experiments reported in Figure 5, and 3 for the experiments reported in Figure 6 and 7.

### Staining and rootlets imaging

Staining analyses (hematoxylin) were performed over two of the extracted rootlets to evaluate the heterogeneity of the sampled fiber population (Supplementary Figure 1). Total fiber count and estimation of diameters were performed using ImageJ64. Contour of each axons were drawn using the freehand selection tool. Areas each contours were measured by the software. Each axon diameter was calculated from the measured area, approximated as a perfect disk.

### Nerve-on-a-chip platform

A nerve-on-a-chip platform was used to record and stimulate the extracted rat dorsal rootlets, as described in the work by Gribi and colleagues^33^. The platform consisted of 2 stimulation electrodes and 8 recording electrodes arranged in a microchannel (Figure 3a). The entire device measured 4 × 2.5 cm^2^. The PDMS microchannels had a cross-section of 100 × 100 µm^2^. Recording and stimulating electrodes were encapsulated in PDMS microchannels (10 mm long). Recording electrodes had contact areas measuring 100 × 300 μm^2^, while stimulation contacts were larger, measuring 100 × 600 μm^2^. Wires were soldered to the chip for connection to an external stimulation (IZ2H stimulator controlled via an RZ2 processor, Tucker Davis Technologies) and recording device (AM system differential AC amplifier, bioamp processor, Sequim, USA and an RZ2, Tucker Davis Technologies). To insert the rootlets into the platform, a suture was tied to one end of each rootlet, allowing it to be gently pulled through the microchannel. All recordings were performed with the nerve-on-a-chip platform located in a faraday cage at room temperature (∼23 °C), using the recording electrode number 6 to maximize the signal to noise ratio^33^, and a sampling rate of 48,828 Hz.

### Data filtering

All data were filtered using a 6th-order, band-pass, digital Butterworth filter with cut-off frequencies of 60-2000 Hz. This range was chosen to remove both the power line noise and the stimulation artefacts. Indeed, since all the tested stimulation conditions used a pulse-width of 50 μs, the duration of the stimulation artefacts was sufficiently short to be efficiently removed without affecting the neural signals (Figure 3b).

### Experiments using single stimulation pulses, BioS bursts, or constant-amplitude bursts

When evaluating the effects of single stimulation pulses, BioS bursts, or constant-amplitude bursts (Figures 3, 4, and 5), repetition frequency of 0.5 Hz were used, to ensure that consecutive neural responses were not affected by the previous stimulation. Each tested stimulation condition was recorded for a minimum of 40 s (i.e., n ≥ 20 repetitions).

To define the minimum stimulation amplitude used for designing a BioS bursts, we searched for the lowest current necessary to induce an observable neural response with a single stimulation pulse. The exact value was different for each rootlet, but remained in the range of 10-15 μA. Similarly, to define the maximum amplitude, we increased the stimulation current until further increments did not elicit observable increases in the neural response. Across the tested rootlets, the maximum current remained in the range of 30-40 μA. When different maximum BioS burst amplitudes where tested (Figure 5), these were set to values within this range. Single stimulation pulses and constant-amplitudes bursts were always delivered at the maximum amplitudes used for the BioS bursts.

Neural responses were extracted using a time-window starting 5 ms before each stimulation pulse/burst and lasting 30 ms. As in the case of the simulated signals, to analyze the induced responses we computed the mean absolute value, the peak to peak amplitude, and the mean standard deviation (Supplementary Figure 5). Mean absolute value and peak to peak amplitudes were computed over the whole extracted signals (30 ms long), for all the stimulation conditions. Instead, the mean standard deviation was only computed over the actual duration of the induced response: 0 to 5 ms after the stimulation onset, in the case of a single pulse; or 0 to 25 ms in the case of a burst. To allow comparisons across different rootlets, each statistic was normalized with the median value obtained with a single pulse of stimulation.

In the case of constant-amplitude bursts (Figure 4), the induced neural activity was also categorized as either an initial response (−1 to 5 ms from the stimulation onset) or subsequent responses (5 to 25 ms from the stimulation onset). Both types of responses were analyzed similarly to the full responses. However, instead of the mean absolute value, in these analyses we computed the sum of the absolute values to evaluate the contribution of these two types of responses to the overall activity.

When assessing the effect of amplitude modulation, we bootstrapped (n iterations = 10,000) the modulation depth of each statistic. This was defined as the difference between the highest and lowest median statistics obtained over the tested amplitudes.

### Experiments using continuous BioS or pulsed stimulation

Single neural responses induced during continuous BioS or pulsed stimulation (Figures 6 and 7) were extracted using time-windows starting at the onset of each BioS burst or stimulation pulse and lasting an entire stimulation period. The extracted responses were analyzed as previously described. Each stimulation condition was tested for 10 s.

Temporal analysis of the mean absolute value (Figure 6b) was performed by grouping the induced neural responses into 2 s windows. The bootstrapped modulation over time (n iterations = 10,000, Figure 6c) was computed as the difference between the medians of the mean absolute values of the neural responses induced during the last 2 and first 2 s of recordings.

### Limitations of the metrics used for analyzing neural responses

The statistical metrics used to probe the synchronicity, level of activity and variability of the induced neural activity may sometimes fail to accurately capture specific aspects of the responses. Anticipating such “failure modes” may help interpret these metrics more accurately (e.g. analogously to how familiarity with the shortcomings of the mean, median and mode helps in the interpretation of these values). As described above, the overall level of activity in the target population was estimated using the mean absolute value (MAV) of the response. In a simple scenario (i.e. no interference), the MAV obtained with 10 individual action potential, and the MAV obtained with a compound action potential made up of 10 fibers would be the same. As such, measuring the MAV over a given time window is a direct measure of the total number of action potentials produced within that time period. However, when action potentials occur in quick succession, the likelihood of interference increases (where the positive and negative phases of an action potential may partially or completely cancel each other out). This effect becomes more likely when a large number of fibers are recruited asynchronously in a short period of time (such as when using high BioS repeating frequencies). Under these conditions, MAV values would tend to underestimate the level of activity induced by stimulation.

Synchronicity was gauged using the peak to peak amplitude of the response. This metric directly captures the height of the largest compound action potential, which is directly proportional to the number of fibers firing synchronously. High values of peak to peak amplitude can only be obtained when a large number of action potentials occur near simultaneously. Indeed, even small temporal shifts would impact the size of the compound action potential measured in the aggregate response. However, similarly to MAV, peak to peak amplitude is susceptible to interference effects. Indeed, two large synchronous responses could partially or totally cancel each other out, leading to an underestimation of the synchronicity of the response. Nevertheless, this is unlikely to have played a meaningful role in our results, since the timing required to cause interference prevents a fiber from interfering with itself, therefore making it difficult for any large compound action potential to be cancelled out (i.e. if two thirds of the fibers are involved in an action potential, only the other third could play a role in any interference effect). However, when measuring highly asynchronous responses, it is plausible that interference may have led to underestimation of the level of synchronicity. In addition, an accurate comparison between the synchronicity of different responses using the peak to peak amplitude is only possible when a similar number of fibers is recruited between the tested conditions. As demonstrated by our computational model (Figure 2b) and ex-vivo experiments (Figure 4), 1000/2000 Hz BioS and constant-amplitude stimulation bursts recruited a similar number of fibers as those recruited by a single pulse of stimulation delivered at the burst maximum amplitude. Therefore, with the exception of when 8000 Hz was used, this condition was largely met.

Finally, the mean standard deviation was used to estimate the variability of neural responses, and therefore the level of stochasticity in the activity. Any changes in the response from one stimulation pulse to the next are captured by this metric. In this case, the most likely effect that may have played a role in our results is the presence of any additional source of variability that may not be associated with changes in neural response due to the stimulation (e.g. temperature fluctuation affecting neural dynamics). Since we assume this type of “external” variability affects all measurements equally, we expect that this effect only accounted for a very small proportion of the measured variability, and therefore did not play any substantial role in the interpretation of our results.

### Statistics and data analysis

Data acquisition and analysis were not performed blind to the experimental conditions. To avoid contaminating the data with sporadic movement artefacts during continuous stimulation experiments, statistics that were larger or lower than the median of the sampled distribution by 4 times the standard deviation were excluded from the analysis. No more than 3.4% of the recorded samples were excluded per trial. No additional data were excluded from the analyses. Statistical significance was analysed using the two-sided, paired, Wilcoxon signed rank test, the two-sided Wilcoxon rank sum test, or a bootstrap test based on the Monte Carlo algorithm resampling scheme (n iterations = 10,000). In all cases, Bonferroni corrections were used to correct for multiple comparisons. No assumptions on data distributions were performed. Additional details about the number of repetitions and the tests used for each experiment are reported in the corresponding figure legends.

### Code and data availability

Recorded data, software routines used for data analysis, and the computational model are available from the corresponding authors upon reasonable request.

## Acknowledgments

We thank Dr. Marco Capogrosso for the fruitful discussions regarding several aspects of the presented work. We would also like to thank Jessica Sordet-Dessimoz (EPFL Histology Core Facility) for her help in histology.

## Author contribution

E.F., E.D. conceived the study, performed the computer simulations and experiments, analyzed the data, and wrote the manuscript. S.G. performed the surgical procedures and performed the experiments. S.M. and S.L. supervised the work. All authors contributed to the editing of the manuscript.

## Competing interests

The authors declare no competing financial interests.

## Supplementary Figures

**Supplementary Figure 1.**
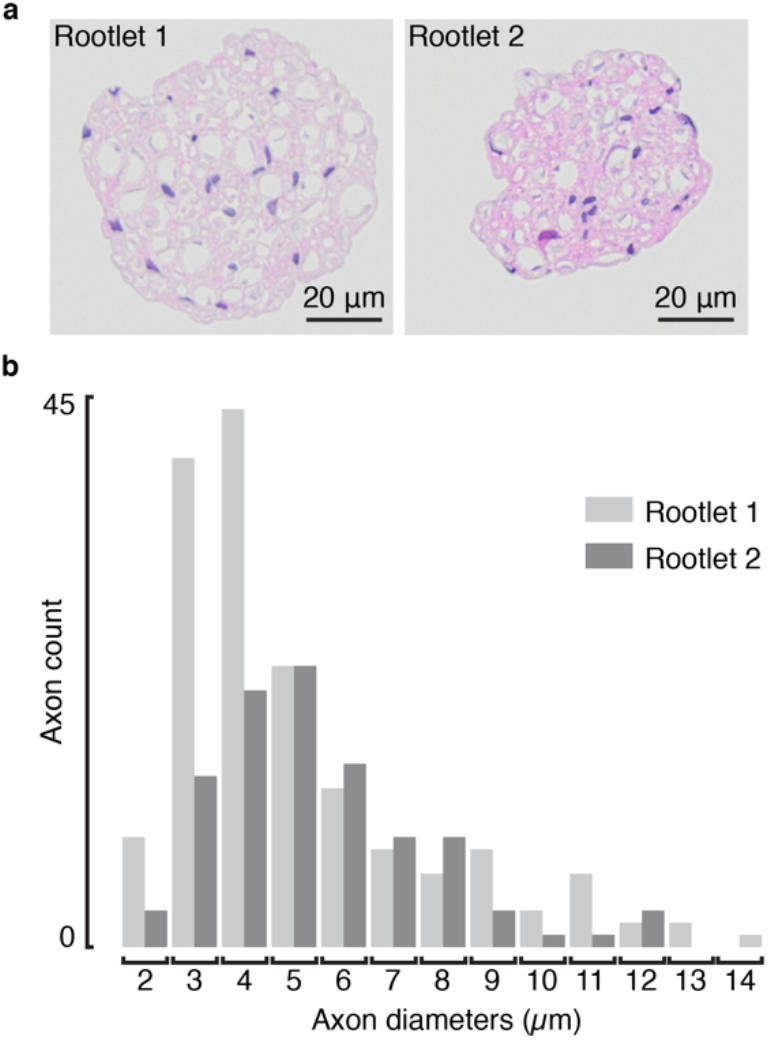
Histological analysis of the extracted dorsal rootlets. **a**, Cross section of Rootlet #1 (left) and #2 (right) used for the experiments reported in Figure 3 and 4. Staining: hematoxylin. **b**, Histogram reporting the number and the diameter of the imaged axons, for the two analyzed rootlets.

**Supplementary Figure 2.**
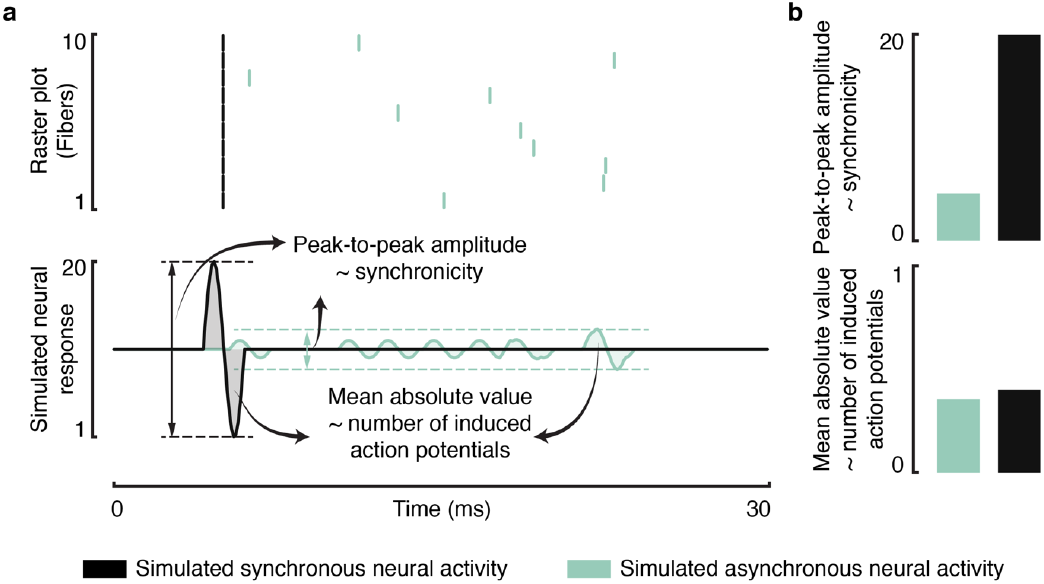
Measuring the synchronicity and number of induced action potentials from recorded neural responses. **a**, Raster plots (top) and simulated neural responses (bottom) generated by synchronous neural activity (simulated as 10 fibers firing all at the same time; black) and asynchronous neural activity (simulated as 10 fibers firing randomly between 0 and 30 ms, following a uniform distribution; aquamarine). Simulated neural responses are computed by convolving fibers action potentials with a sinusoidal waveform of 2 ms duration. **b**, Bar plots representing the peak-to-peak amplitude and mean absolute value of the simulated neural responses, statistics that we used as proxy to measure the synchronicity and number of action potentials, respectively. As expected, the peak-to-peak amplitude is greater for synchronous neural activity compared with asynchronous neural activity (since summation of multiple synchronous action potentials causes larger neural responses), while the mean absolute values are similar (since the same number of action potentials were simulated).

**Supplementary Figure 3.**
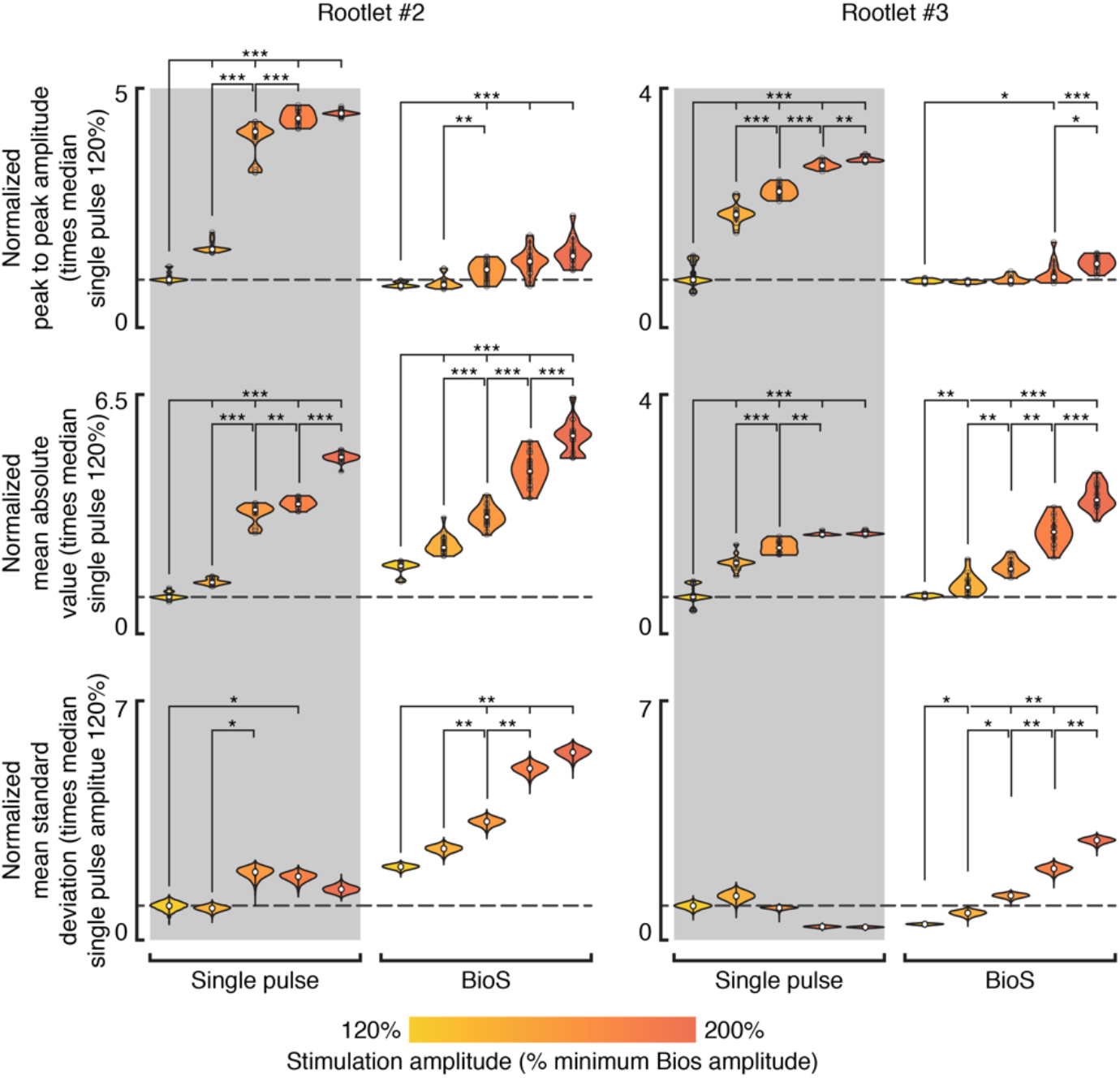
Extended data of Figure 5. Violin plots reporting, from top to bottom, the peak to peak amplitude, mean absolute value, and mean standard deviation of the neural responses induced by the tested stimulation conditions, for rootlet #2 (n = 260 responses) and #3 (n = 454 responses). Each statistic is normalized with respect to the median value of single pulse stimulation, as highlighted by the dashed lines. The violin plots for the peak to peak amplitude and the mean absolute value report the sampled distributions. Violin plots for the mean standard deviation report the bootstrapped distribution (n iterations = 10,000). Stars indicate a significant difference: ***, p<0.0001; **, p<0.005; two-sided Wilcoxon rank sum test with Bonferroni correction for multiple comparisons.

**Supplementary Figure 4.**
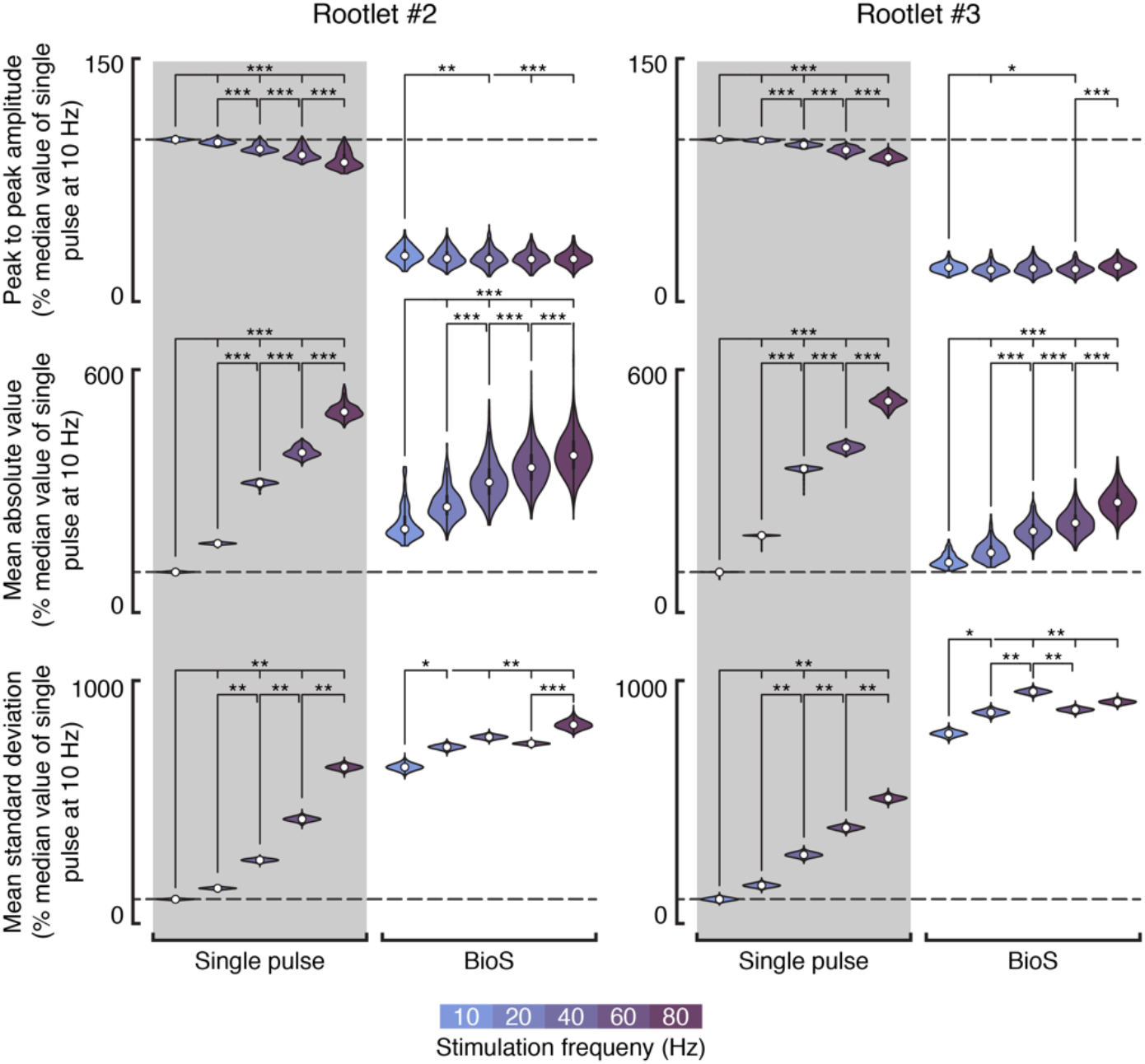
Extended data of Figure 7. Violin plots reporting, from top to bottom, the peak to peak amplitude, mean absolute value, and mean standard deviation of the neural responses induced by the tested stimulation conditions, for rootlet #2 (n = 57,737 responses) and #3 (n = 58,183 responses). Each statistic is normalized with respect to the median value of single pulse stimulation, as highlighted by the dashed lines. The violin plots for the peak to peak amplitude and the mean absolute value report the sampled distributions. Violin plots for the mean standard deviation report the bootstrapped distribution (n iterations = 10,000). Stars indicate a significant difference: ***, p<0.0001; **, p<0.005; two-sided Wilcoxon rank sum test with Bonferroni correction for multiple comparisons.

**Supplementary Figure 5.**
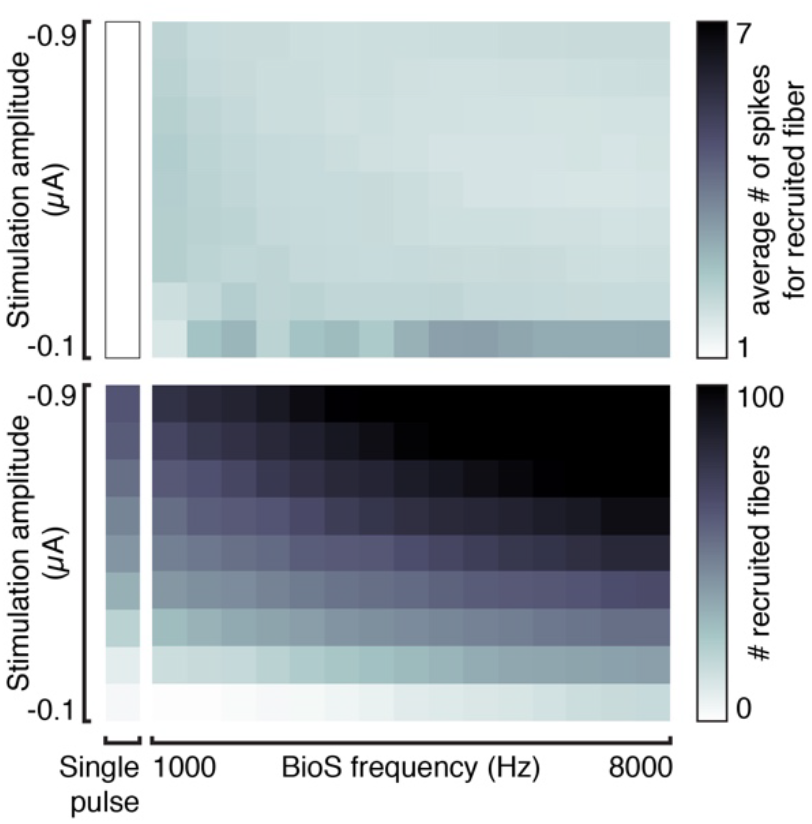
BioS characterization at room temperature. Effect of BioS burst frequency and maximum amplitude on the number of action potentials (APs) generated in each fiber recruited by the stimulation (top), and on the number of recruited fibers itself (bottom). Simulations performed at 23° C to match the experimental conditions of the recordings performed with the nerve-on-a-chip platform.

